# Supplementation via DAF-16 and *pnk-1* driven pantothenate–coenzyme A flux improves disease related stress resistance in *C. elegans*

**DOI:** 10.64898/2026.07.08.737381

**Authors:** Steffi M Jonk, Alan Nicol, Alex Braun, Wenry Wang, James R Tribble, Peter Swoboda, Pete A Williams

## Abstract

Metabolic pathways are increasingly recognized as tractable targets in aging and disease. Building on prior work demonstrating that supplementation with low–molecular weight metabolites (amino acids, vitamins, and their intermediates) can extend lifespan in *Caenorhabditis elegans*, we focused on pantothenate (vitamin B_5_), which is dysregulated in sarcopenic muscle and in several neurodegenerative and metabolic disorders. Pantothenate is the obligate precursor of coenzyme A through a short, highly conserved biosynthetic pathway in which loss-of-function mutations can cause neurodegeneration with brain iron accumulation. In *C. elegans*, the longevity curtailing transcription factor DAF-16/FOXO has a conserved binding element in the promoter region of *pnk-1*, encoding the first enzyme (PNK-1) in the coenzyme A pathway, and *pnk-1* is markedly upregulated in long-lived *daf-2* (insulin/-like receptor) mutants, implicating coenzyme A metabolism in longevity. Here, we demonstrate that CoA levels naturally increase during early life and decrease towards older age in *C. elegans*. Dietary pantothenate supplementation increases coenzyme A levels with minimal effects on lifespan but systemic effects on lipid metabolism, mitochondrial dynamics, and muscle structure under basal conditions. Under DAF-16–associated stress conditions, including heat and oxidative stress, *pnk-1* expression is upregulated and pantothenate supplementation robustly extends lifespan and improves mobility. Finally, we demonstrate dysregulation of *daf-16* and *pnk-1* expression in amyotrophic lateral sclerosis (ALS) models, in which pantothenate supplementation confers both lifespan extension and cholinergic neuroprotection.

## INTRODUCTION

Understanding nutrition and metabolome in relation to health and disease has provided insight into nutrients in cancer spread (Abbott et al., 2026), metabolic dysfunction in neurodegenerative disease (Tribble et al., 2025), and using metabolomics facilitates estimating biological age and mortality risk (Anagnostakis et al., 2025). Aiming to understand how the changing metabolome influences aging and neuronal health, we recently demonstrated metabolic changes in sarcopenic muscle that directly affect neuronal and neuronal metabolic health. Empirical testing of candidate sarcopenic muscle metabolites in *C. elegans* revealed that supplementation of some of these identified metabolites extended lifespan in *C. elegans* and provided metabo- and gero-protection (Jonk et al., 2025). We identified dysregulated muscle pantothenate (PA) with sarcopenia and age, in support of other studies carried out in rodents (Jonk et al., 2025; Uchitomi et al., 2019). Altered levels of PA have been demonstrated in amyotrophic lateral sclerosis (ALS) (Barros et al., 2023), Alzheimer’s disease (Xu et al., 2020) and type II diabetes (Shoura et al., 2025) patients suggesting it might have a more general role in metabolic health.

PA is the essential B-vitamin (B_5_) that is synthesized to coenzyme A (CoA) via a short and well-conserved pathway identified in eukaryotes, plants and archaea; while bacteria are identified with a different CoA pathway (Leonardi & Jackowski, 2007; Webb et al., 2004). CoA is essential in humans and animals for the anabolism of lipids and heme, the catabolism related to the TCA cycle, acetylation with a signaling- and gene expression output, fatty acid synthesis, and folate metabolism. Mutations in the genes implicated in the CoA pathway can be lethal or result in disease (Barritt et al., 2024). The CoA pathway initiates with pantothenate kinase (*PANK*)that phosphorylates PA to 4’-phoshopantothenate, *PPCS* condenses this with a cysteine group to 4’-phosphopantothenoylcysteine, *PPCDC* decarboxylates this to 4’-phosphopantetheine, finalized by an addition of AMP from ATP to form dephospho-CoA and phosphorylation to CoA by *COASY* (Leonardi et al., 2005) (**Figure 1A**). Mutations in *PANK2* (mostly expressed in the nervous system) and *COASY* result in *PKAN* and *CoPAN*; *PANK*-or *COASY-*associated neurodegeneration, which are degenerative disorders characterized by neurodegeneration with brain iron accumulation (NBIA) (Hayflick et al., 2018). Exploring the therapeutic effects of supplementing PA or derivatives and altering expression levels of *PANK4* demonstrated favorable lipid- and glucose metabolism (Miranda-Cervantes et al., 2025; Rumberger et al., 2011).

**Figure 1.**
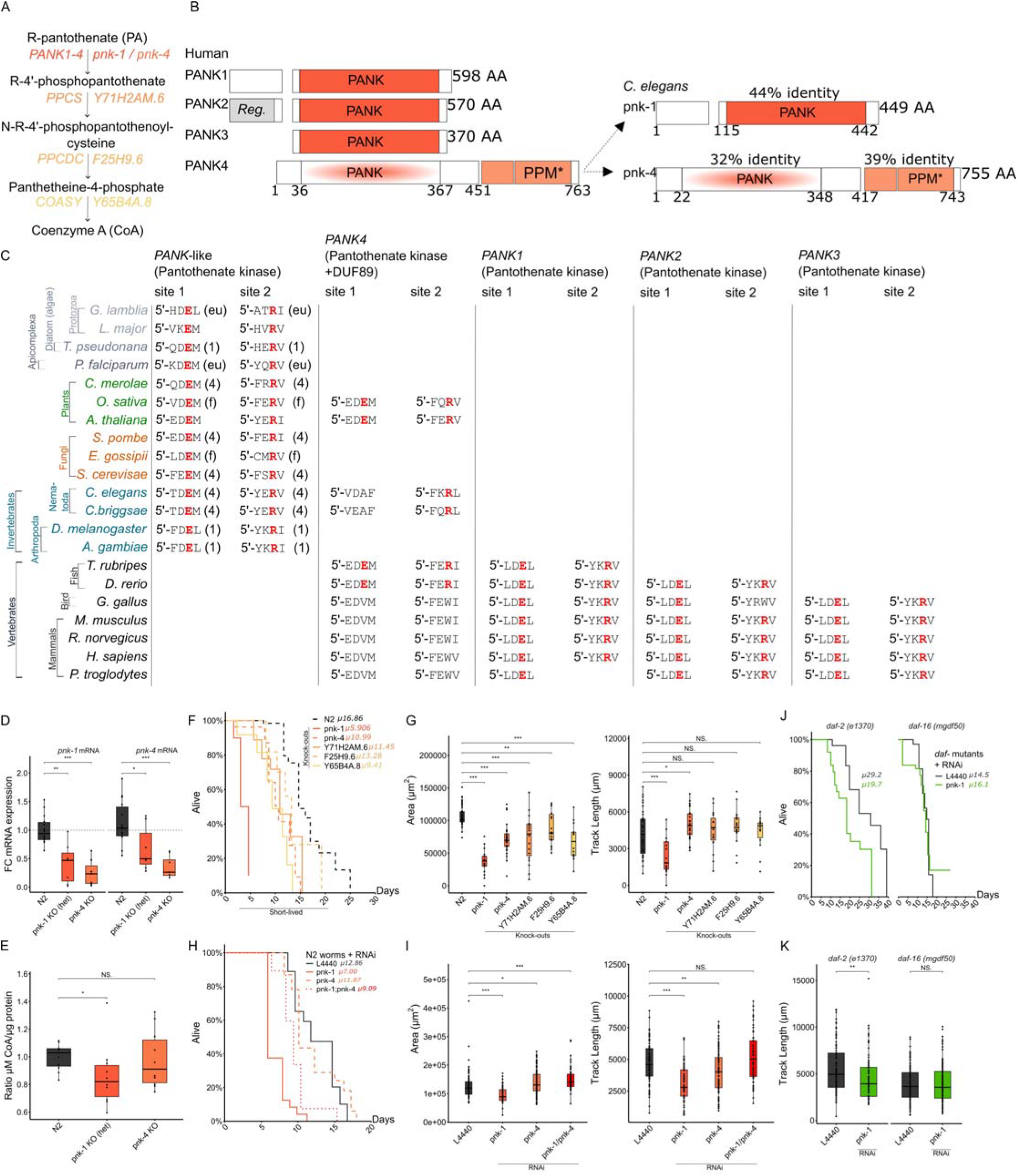
Pantothenate kinases are highly conserved, and *pnk-1* is essential for *C. elegans* survival and regulated via *daf-16*/FOXO. (**A**) Representation of the PA—CoA pathway with human genes on the left and *C. elegans* genes on the right. (**B**) Representation of PANK proteins in human and *C. elegans*, based on Miranda-Cervantes et al., Nature Comm. 2025 (Miranda-Cervantes et al., 2025). (**C**) Functional alignment of PANK proteins in several organisms, demonstrating PANK-like proteins, PANK4, PANK1, PANK2 and PANK3. (**D**) mRNA expression and (**E**) CoA levels of *pnk-1* and *pnk-4* knockouts. (**F**) Lifespan by Kaplan-Meier curves of PA-CoA *C. elegans* knockouts, and (**G**) the according growth phenotype and mobility (by track length) as recorded by Wormlab. (**H**) Lifespan by Kaplan-Meier curves of RNAi of *pnk-1, pnk-4* and double *pnk-1;pnk-4* and (**I**) the according growth phenotype and mobility by track length. (**J**) Lifespan by Kaplan-Meier curves of RNAi of *pnk-1, pnk-4* and double *pnk-1;pnk-4* in *daf-2* (*e1370*) and *daf-16* (*mgdf50*) mutants and (**K**) the according mobility by track length. *p = * < 0.05, ** < 0.01, *** < 0.001, **** < 0.0001.

*C.egans* is widely used in longevity research because of its short lifespan, fast-generation time and especially since the discovery of a number of long- and short-lived mutants, prominently including mutations in *daf-2* and *daf-16*, respectively (Kenyon et al., 1993). The transcription factor DAF-16 has a conserved binding element in the promoter region of *pnk-1* (Lee et al., 2003) (**Figure S1A**), the *C. elegans* homolog of human *PANK4*, which encodes one of the two pantothenate kinases identified in *C. elegans* (**Figure 1B**). Additionally, *pnk-1* is five-fold upregulated in *daf-2* mutants and the *daf-2* doubled lifespan shortens with *pnk-1* RNA interference (Lee et al., 2003). Here we explore in *C. elegans* the CoA pathway and the role of *pnk-1* in longevity, healthy aging and stress resistance, and its potential as a therapeutic target in ALS.

## RESULTS AND DISCUSSION

PA is an essential, diet-derived vitamin with poorly characterised bioavailability. Dietary intake of PA is largely unknown (Freese et al., 2023). There are several mechanisms to maintain the pool of PA-CoA: cellular PA pools are maintained via CoA hydrolysis and pantetheine recycling, and uptake occurs via the SLC5A6/SMVT transporter, including across the blood-brain barrier. (Freese et al., 2023). The first step in the PA-CoA pathway in eukaryotes is a rate-determining step of pantothenate kinase (PANK) that phosphorylates PA to 4′-phosphopantothenate (Leonardi et al., 2005). Mammals (including humans) have four *Pank* genes (*-1, -2, -3, -4*) (**Figure 1B**) which are primarily expressed in different tissues (*Hs*; *-1*: Liver, kidney; *-2*: CNS; *-3*: intestine; *-4*: skeletal- and cardiac-muscle (Human Protein Atlas)). Human PANK4 is a pseudo-pantothenate kinase: the dysfunctional kinase domain is missing the functional Glu138 and Arg207 amino acids (Yao et al., 2019) (**Figure 1C**). PANK proteins are implicated in disease; from ulcerative colitis, where upregulation of PANK3 reconstructed the intestinal barrier and folic acid (vitamin B_9_) was identified as an activator of PANK3 (Zhang et al., 2025), to specific associated phenotypes of mutations in *Pank* genes in mice and human (**Supp. Table 1**). These phenotypes include decreased fatty acid oxidation, abnormal gluconeogenesis, increased body weight (*Pank1*), male infertility and small testis (*Pank2*), decreased locomotor activity (*Pank3*), cataract and hyperactivity (*Pank4*). Together these data support a clear role for PA and the PANK family of genes/proteins across a wide range of metabolic functions.

Two pantothenate kinase genes are present in *C. elegans* (*pnk-1* and *pnk-4*), which are both orthologous to the human *Pank4* gene with 67% and 59% protein homology (AlphaFold) (**Figure 1B, Supp Table 3**). To better understand the functions and roles of PANK proteins, we explored protein alignments across species. Cross-species alignments revealed a clear evolutionary trajectory: organisms with a single PANK encode a fumble-domain kinase with a separate ARMT-1 phosphatase (*e*.*g. D. melanogaster;* the *Fbl (fumble)* gene, is the gene in *melanogaster* that encodes the pantothenate kinase protein, mutants demonstrate severe movement characteristics (Afshar et al., 2001)); organisms with two PANK proteins add a PANK4-like kinase/phosphatase fusion (*e*.*g. C. elegans*); and vertebrates have elaborated this into four paralogues (PANK1–4) through gene duplication, with progressive loss of PANK4 kinase activity in birds and mammals (**Figure 1C, Supp. Dataset 1**).

To characterize the functional requirement for PNK-1 and PNK-4 in *C. elegans*, we assessed knockout and RNAi phenotypes across the entire PA–CoA pathway in *C. elegans*, which is conserved to the human pathway given that it has 4 enzymatic steps from PA to CoA (**Figure 1A**), and protein homology ranges between 44% and 68% with conserved protein domains (**Supp. Table 3**). Knockout (KO) worms of the implicated genes in the PA-CoA pathway are available (Au et al., 2019; Consortium, 2012) (for details of the deletions in the coding regions of the respective gene see **Supp. Table 5**). It was previously described that the *pnk-1* KO strain was not viable and CoA supplementation partially rescued the detrimental phenotype (Srinivasan et al., 2015). This was also observed in *Drosophila melanogaster*, highlighting the importance and conservation of this pathway. RT-qPCR of *pnk-1* and *-4* confirmed decreased expression in KO mutants, leading to decreased CoA levels as determined by luminescence (**Figure 1D, E**). We confirmed the drastic lifespan decrease of *pnk-1* KO mutants and demonstrated decreased lifespan and altered growth phenotypes (worm size) and mobility (track length) compared to wild-type N2 of *pnk-4, Y71H2AM*.*6, F25H9*.*6* and *Y65B4A*.*8* KO mutants (**Figure 1F, G**). The viability and unchanged CoA levels of *pnk-4* KO worms indicate that, analogous to human *Pank4, pnk-4* does not function as a pantothenate kinase. To understand and confirm the detrimental effects of reduced PNK-1 and -4 expression we performed RNAi of *pnk-1, pnk-4*, and double *pnk-1;pnk-4*. Lifespan and mobility decreased drastically for *pnk-1* and *pnk-1;pnk-4*, and slightly for *pnk-4* (**Figure 1H, I**)

DAF-16, the transcription factor (TF) known for its role as stress-associated and -activated and required for the doubled lifespan extension in *daf-2* mutants, has a binding element (DBE, **Figure S1A**) in the promoter sequence of *pnk-1* (384 bp upstream). This DBE in *pnk-1* is conserved in *Caenorhabditis* species and identified in mouse and human *Pank1* and *-2* proteins (**Supp. Table 4**). Long-lived *daf-2* mutants have 5-fold upregulated *pnk-1* expression (Lee et al., 2003) and RNA interference (RNAi) of *pnk-1* in the long-lived *daf-2* mutants decreased their lifespan (Lee et al., 2003). To confirm the detrimental effects of *pnk-1* RNAi and to validate that *pnk-1* is regulated via *daf-16*, we performed RNAi of *pnk-1* in *daf-2* (*e1370*) and *daf-16* (*mgdf50*) mutants, which demonstrated that *pnk-1* RNAi had an effect in *daf-2* mutants by decreased lifespan and mobility, but not in *daf-16* mutants (**Figure 1J, K**). Together these support that *pnk-1* is essential in worms and regulated via DAF-16.

We next assessed the temporal dynamics of CoA levels in N2 worms which demonstrated increased CoA output in young adults (day 1 to 5), followed by a decline in aged adults (toward day 10 to 15) (**Figure 2A**). Increased PA levels have been demonstrated in aged mouse skeletal muscle (Jonk et al., 2025; Uchitomi et al., 2019), which could be explained by the decreased CoA output later in life that was observed in *C. elegans*. It must be demonstrated whether this is true in humans and, or mouse. Our previous data demonstrated that PA supplementation increased lifespan in *C. elegans* (supplemented via NGM) (1). Worms supplemented with 5 mM PA (via dead *E. coli* OP50), demonstrated increased CoA levels after PA supplementation (**Figure 2B**), however, no overt lifespan extending or mobility effects in wild-type worms were observed after direct PA or CoA supplementation (every 3 days) (**Figure 2C**); which could be explained by the difference in method of supplement administration.

**Figure 2.**
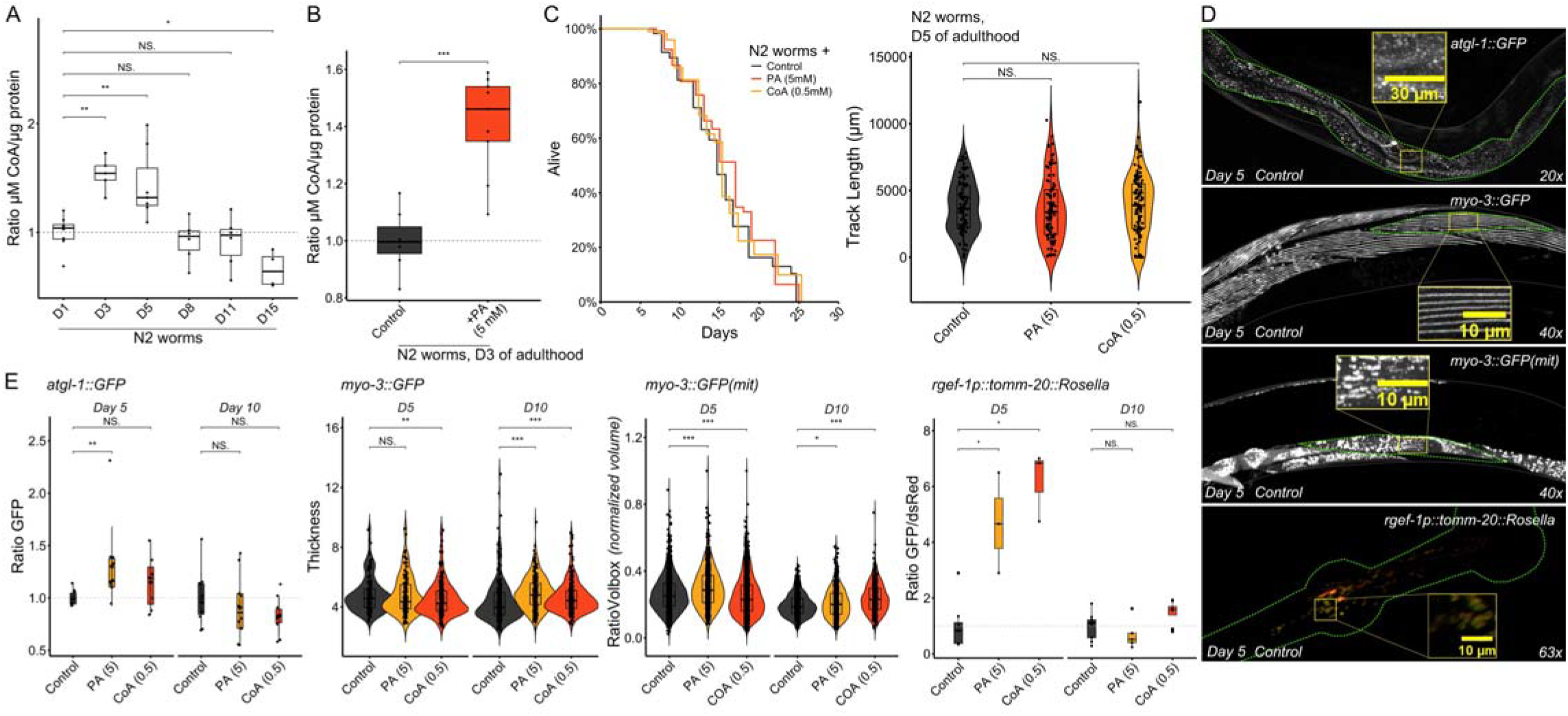
Supplementation of PA and CoA has widespread physiological effects. (**A**) CoA levels during *C. elegans* aging at day 1, 3, 5, 8, 11 and 15 of adulthood. (**B**) CoA levels after PA (5 mM) supplementation from day 1 to day 3 of adulthood (measured at day 3 of adulthood). (**C**) Lifespan by Kaplan-Meier and mobility by track length of N2 worms supplemented with PA (5 mM) or CoA (0.5 mM). (**D, E**) Reporter strains presenting atgl-1::GFP (VS20), myo-2::GFP (RW1596), myo-2::GFP(mit) (SJ4103) and rgef-1p::tomm-20::Rosella (SJZ42), and the respective quantified ratio of GFP intensity, thickness, RatioVolBox (captures how “space⍰filling” a mitochondrion is inside its bounding box; high ratiovolbox: the mitochondrion nearly fills its bounding box, typical of round, swollen, or compact mitochondria. Low ratiovolbox; the mitochondrion occupies only a small fraction of the bounding box, typical of elongated, branched, filamentous mitochondria) and ratio of GFP/dsRED intensity, at day 5 and 10 of adulthood after PA (5 mM) and CoA (0.5 mM) supplementation. *p = * < 0.05, ** < 0.01, *** < 0.001, **** < 0.0001.

To investigate systemic effects of PA and CoA supplementation at a tissue-specific level, we used reporter strains and microscopy for *atgl-1 (Zhang et al., 2010)* (a rate-limiting lipolytic enzyme, demonstrated to increase during fasting (Zaarur et al., 2019)), myosin (Campagnola et al., 2002), mitochondria and mitophagy (Cummins et al., 2019) (**Figure 2D**). These reporter strains demonstrate widespread physiology by lipolysis in the intestine (by *atgl-1*; adipose triglyceride lipase), heavy chain myosin filaments in skeletal muscle (by *myo-3p*; essential for body contraction), mitochondrial phenotype in skeletal muscle (by *myo-3p::mito*, targeting only the mitochondria in skeletal muscle), and by mitophagy in neurons by GFP quenching in low pH environments (like lysosomes under *rgef-1p*; Ras guanyl-releasing protein 3 expressed in the nervous system). *atgl-1* expression by GFP intensity increased at day 5 of adulthood, but not at day 10, following PA supplementation (**Figure 2E, S2C-D**). Like *atgl-1*, mitophagy by fluorescent intensity (Ratio of GFP/dsRED) changed at day 5 of adulthood, but not at day 10 (**Figure 2E**). PA and CoA supplementation improved myosin organization and structure at day 10 of adulthood, suggesting a better state of striated muscle fibres. Mitochondrial morphology by sphericity and normalized volume (ratiovolbox) demonstrated decreased volume and sphericity at day 10 of adulthood compared to day 5 (**Figure S2C**); suggesting fragmentation or irregularity by *e*.*g*. increased fission and decreased fusion. Again, supplementation with PA increased ratiovolbox and sphericity at day 5 of adulthood, while CoA supplementation demonstrated this at day 10 of adulthood (**Figure 2E, S2C**). PA and CoA supplementation reduced the mitophagy ratio (GFP/dsRED) in neurons at day 5 (**Figure 2E, S2D**), consistent with a lower burden of damaged mitochondria under supplementation. By day 10, the baseline mitophagy ratio was reduced relative to day 5 (**Figure S2C**), reflecting the expected age-associated increase in mitophagic flux. The peak in CoA output at day 5 likely explains the preferential efficacy of PA supplementation at this timepoint: supplementation may be most effective when endogenous *pnk-1*-driven CoA biosynthesis is active. We observed effects in lipid metabolism in the intestine, myosin/muscle, mitochondria and mitophagy in neurons after PA and CoA supplementation, suggesting that in *C. elegans* this pathway is not tissue or organ specific.

To further understand the regulation of the PA-CoA pathway, we assessed mRNA expression and CoA levels in long-lived mutants, which demonstrated decreased *pnk-1* expression in *daf-16* (*mgfd50)*, but an increase of CoA levels, confirming altered CoA regulation in this *daf-16* mutant (**Figure 3A, B**). PA and CoA supplementation to *daf-2 (e1370), daf-16 (mgdf50)* and *daf-2::daf-16 (m65::mgdf50)* mutants showed no effects on longevity, but increased mobility in *daf-2* mutants; confirming that these *daf-2* and *daf-16* mutations affect *pnk-1* expression and this expression dictates the response to PA supplementation (**Figure 3C**). Since DAF-16 is a stress associated TF, we next questioned whether PA supplementation had effects under DAF-16 associated stress, which include heat-stress (Henderson & Johnson, 2001), oxidative stress and hypoxic stress (Leiser et al., 2013).

**Figure 3.**
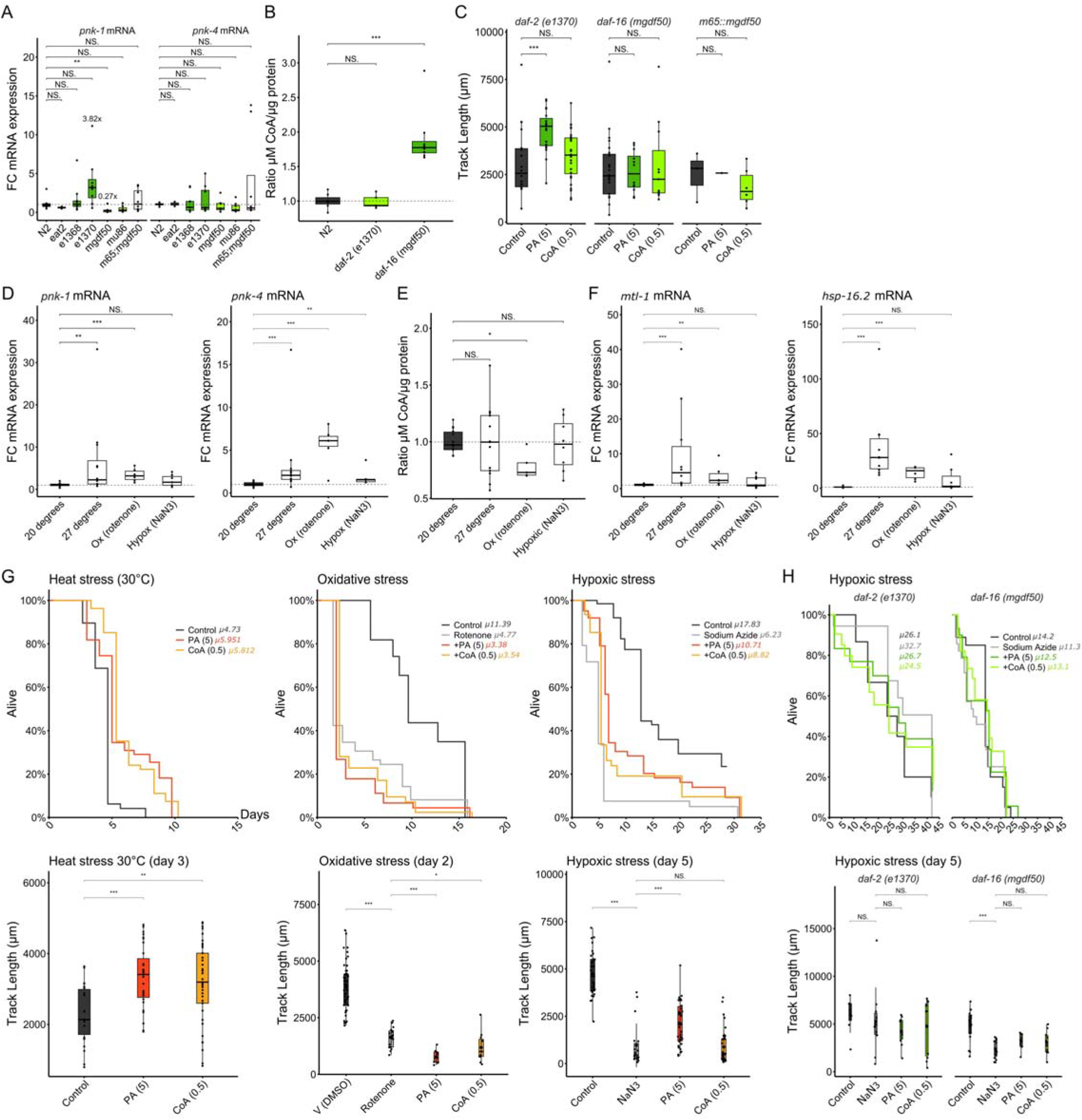
Supplementation of PA and CoA protects against DAF-16 associated stressors. (**A**) mRNA expression and (**B**) CoA levels in long-lived and *daf-* mutants. (**C**) Mobility by track length of *daf-* mutant worms supplemented with PA (5 mM) or CoA (0.5 mM). (**D)** *pnk-1* and *pnk-4* mRNA levels by RT-qPCR and (**E**) CoA levels in N2 worms exposed to heat stress at 30° C, rotenone (1.5 µM) or sodium azide (2 mM) from day 1 to 3 of adulthood (measured at day 3). (**F**) *mtl-1* and *hsp-16*.*2* mRNA expression of N2 worms exposed to heat, oxidative and hypoxic stress. (**G**) Lifespan by Kaplan-Meier and mobility by track length of wild-type N2 worms exposed to heat stress at 30° C, rotenone (1.5 µM), or hypoxic stress by sodium azide at 2 (day 1-2) and 4 mM (day 2 till end of life). (**H**) Lifespan by Kaplan-Meier of *daf-2* (*e1370*) and *daf-16* (*mgdf50*) mutant worms exposed to hypoxic stress by sodium azide at 2 (day 1-2) and 4 mM (day 2 till end of life). *p = * < 0.05, ** < 0.01, *** < 0.001, **** < 0.0001.

We first confirmed altered CoA metabolism during heat stress (30° C), oxidative stress (rotenone, a mitochondrial complex I inhibitor) and hypoxic stress (sodium azide, NaN_3_). We observed increased *pnk-1* after heat- and oxidative stress and increased *pnk-4* expression after heat-, oxidative- and hypoxic stress (**Figure 3D**). CoA output was only significantly decreased after oxidative stress exposure, but not after heat- and hypoxic stress, however some variability was observed suggesting that this pathway, like other metabolic pathways, is possibly sensitive to *e*.*g*. environmental changes. However, the dysregulated PA-CoA metabolism was regulated via *daf-16* by increased expression of the downstream targets *mtl-1* (metallothionein) and *hsp-16*.*2* (heat-shock protein) (**Figure 3H**) and a direct increase of *daf-16* mRNA (**Figure S3D**). However, DAF-16 mRNA expression does not necessarily mean activation and translocation of DAF-16, we therefore explored whether DAF-16 translocated to the nucleus upon stress exposure, which was confirmed for all tested stressors (**Figure S3E**).

We then observed minor effects of CoA supplementation under DAF-16-associated heat stress (30° C) and hypoxic stress (sodium azide) and greater effects of PA supplementation, by a respective increase in lifespan and track length, demonstrating improved mobility (**Figure 3G**). These effects were not observed in *daf-2* and *daf-16* mutants, where hypoxic stress (sodium azide) did not decrease lifespan and mobility in *daf-2* mutants; as previously described (*daf-2* (*e1370*)) demonstrates azide resistance (31)) (**Figure 3H**). Under oxidative stress induced by rotenone, PA and CoA supplementation showed detrimental effects; suggesting that rotenone induced stress did not activate DAF-16 (**Figure 3G**). Although rotenone induced stress activated DAF-16 and downstream targets, PA and CoA supplementation was detrimental. Rotenone inhibits mitochondrial Complex I, generating superoxide and depleting the mitochondrial membrane potential. Under conditions of Complex I inhibition, increased CoA flux may exacerbate mitochondrial substrate overload by driving fatty acid β-oxidation and acetyl-CoA generation in a context where the TCA cycle entry is impaired, thereby worsening rather than resolving the energetic deficit. The detrimental effect of PA and CoA supplementation under rotenone stress therefore likely reflects the pathway-context dependence of CoA metabolism: supplementation is beneficial when DAF-16 is the dominant stress sensor and the electron transport chain is intact, but counterproductive when mitochondrial respiration is directly compromised.

To explore the modulation of pantothenate kinases in longevity and healthy aging, we generated *C. elegans* overexpression (OE) strains. *C. elegans pnk-4*, double *pnk-1::-4*, and *Y65B4A*.*8* (the last enzyme in the PA-CoA pathway) were overexpressed under the Prpl-28 promotor for ubiquitous expression (*rpl-28::cDNA of interest::GFP*) (mRNA expression quantified in **Figure S4A**). It proved impossible to generate overexpression strains for *pnk-1*, which together with the data from the *pnk-1* KO strain (not viable) suggests that pantothenate kinase activity is tightly regulated, and even slight changes in expression may result in detrimental development and growth/lethality. This is line with the observation in *D. melanogaster* where the homolog *dPank* (*Fbl* gene) is required for proper mitosis and meiosis and thereby development (Afshar et al., 2001). CoA levels decreased in *pnk-4* OE strains, while no changes were observed for *pnk-1::-4* and *Y65B4A*.*8* OE strains (**Figure 4A**). Overexpression of *pnk-1::-4* and *pnk-4* had minor effects on lifespan (significant decrease compared to N2 but not to the control strain) under basal conditions, while *pnk-4* OE decreased mobility by track length and worm growth (size) (**Figure S4B**). Under heat stress (27 °C) OE of *pnk-1::-4* and *Y65B4A*.*8* decreased lifespan compared to controls, confirming the increased requirement for CoA during stress (**Figures 4B and 4C**). It was recently demonstrated that overexpression of mouse *PANK4* (using viral vectors) in skeletal muscle had no effect on muscle mass but altered glucose uptake and metabolism: acetyl-CoA levels decreased in skeletal muscle (Miranda-Cervantes et al., 2025). Together, these results from *C. elegans* and mouse demonstrate that CoA metabolism and output is altered in overexpression scenarios and that upregulation of the pathway itself does not protect under basal conditions or during stress.

**Figure 4.**
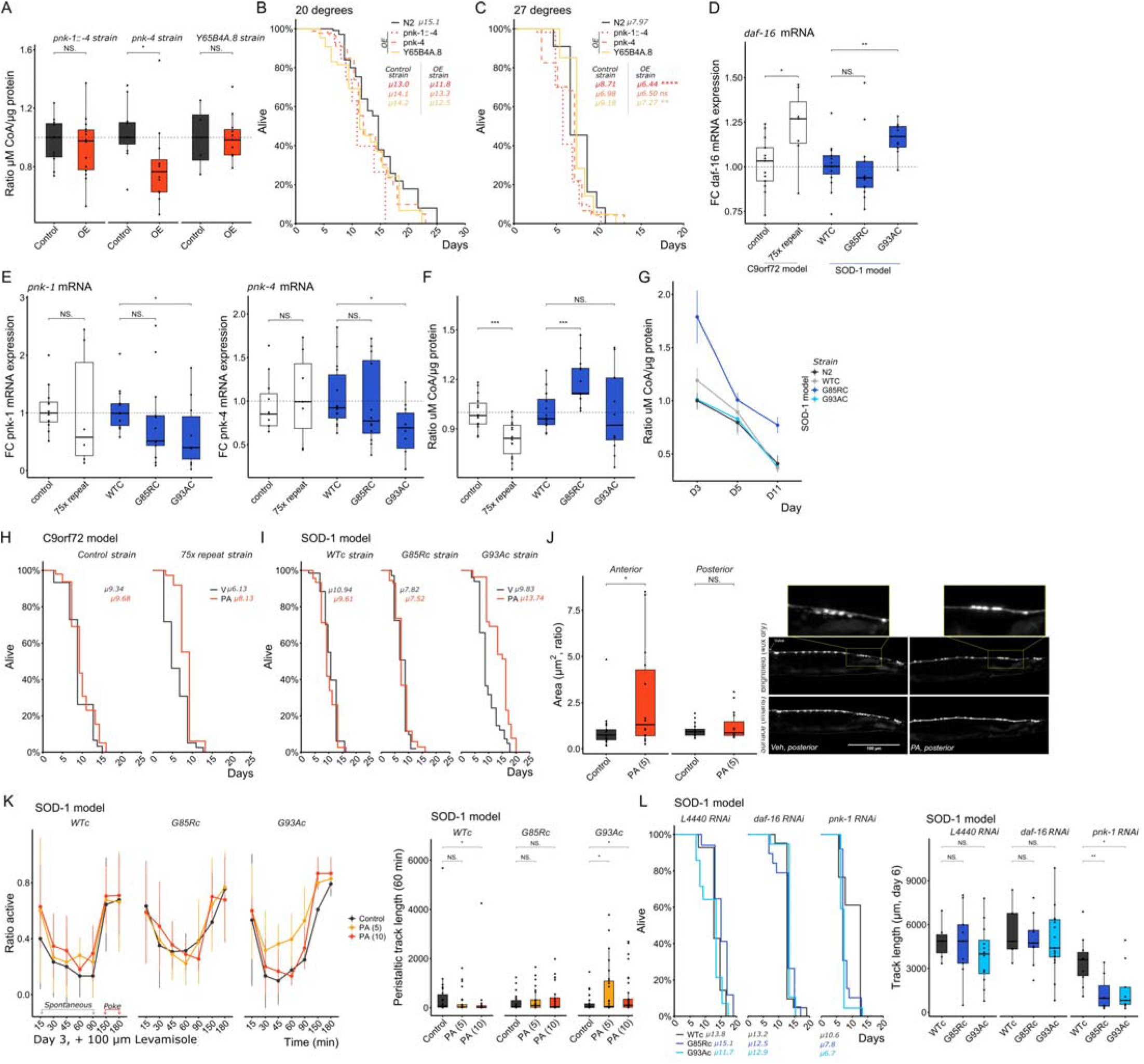
Overexpression of genes in the PA-CoA pathway is detrimental under DAF-16 associated stress and PA supplementation in ALS models alleviates lifespan and mobility phenotypes and provides neuroprotection. (**A**) CoA levels in *pnk-1::-4, pnk-4* and *Y65B4A*.*8* overexpression (OE) strains compared to their respective control. (**B, C**) Lifespan by Kaplan-Meier of *pnk-1::-4, pnk-4* and *Y65B4A*.*8* OE strains at 20 and 27° C. (**D**) *daf-16* mRNA expression levels in *C. elegans* transgenic ALS models of *C9orf72* and *SOD-1*. (**E**) *pnk-1* and *pnk-4* and (**F**) CoA levels in *C. elegans* transgenic ALS models (*C9orf72* and *SOD-1*). (**G**) Temporal assessment of CoA levels in N2 compared to *SOD-1* WTC, G85RC and G93AC strains. (**H, I**) Lifespan by Kaplan-Meier of *C. elegans* transgenic ALS models supplemented with 5 mM PA. (**J**) Neuroprotection shown by the area as a ratio to the control of the reconstructed cholinergic neuron including neuronal bodies and axon of the *C. elegans* transgenic ALS model G93Ac (unc-17::GFP) on day 8 of adulthood, supplemented with 5 mM PA and exposed to 27° C. (**K**) Ratio of spontaneous and poking-induced-activity observed in ALS *SOD-1* worms treated with 5 and 10 mM PA (for 3 days) followed by exposure to 100 µM levamisole (for 3 hours), and mobility by track length at 60 min after levamisole exposure. (**L**) Lifespan by Kaplan-Meier and mobility by track length of ALS *SOD-1* worms exposed to *daf-16* and *pnk-1* RNAi; mobility was measured at day 6 of adulthood. *p = * < 0.05, ** < 0.01, *** < 0.001, **** < 0.0001.

It was previously reported that ALS patients with greater functional decline (lower ALSFRS-R scores) had significantly lower dietary intake of pantothenic acid (*p* = 0.037) (Barros et al., 2023). Our previous research using *C. elegans* models of ALS (Jonk et al., 2025) did not demonstrate beneficial effects of PA. However, to understand whether PA can be used in these *C. elegans* models when DAF-16/FOXO is dysregulated, we used data from the *ALS cell atlas* as a reference (Månberg et al., 2021); which demonstrated increased levels of *Foxo3* (the human ortholog of *daf-16*), but decreased *Pank1* levels in a *SOD-1* G93A mouse model (**Figure S5A**). Since in *C. elegans* a beneficial effect of PA supplementation under DAF-16 activation/translocation was observed, we measured *daf-16, pnk-1* and *-4* mRNA expression in two *C. elegans* ALS models, to further understand the role of *daf-16* and *pnk-1* in different ALS models. We used *Ce* ALS models with an insertion of human wildtype and mutant *C9orf72*; respectively a control and a mutant with 75 GGGCC repeats strain (34) and a model with insertion of wildtype and mutant *SOD-1;* respectively a WTc, mutant G85Rc and G93Ac strain (c representing the method, Crispr/Cas9 (Baskoylu et al., 2018)).

Compared to control/wildtype, we observed increased *daf-16* expression in the *C9orf72* mutant model and the G93Ac mutation in the *SOD-1* model, but not in the G85Rc mutant (**Figure 4D**). *pnk-1* and *-4* expression decreased significantly in worms carrying the G93Ac mutation (**Figure 4E**). However, CoA levels decreased in the *C9orf72* mutant model (respective to the control strain) but increased in the *SOD-1* G85R model (respective to the WTc strain) (**Figure 4F**). To understand natural variation in CoA levels during life, we measured CoA levels in the *SOD-1* strains and compared them to N2 worms at day 3, 5, and 11 of adulthood, which demonstrated increased levels during life for the *SOD-1* G85Rc strain, suggesting genotype specific CoA regulation (**Figure 4G**). We demonstrate altered *daf-16* regulation and *pnk-* and CoA metabolism in two ALS models, but a mutation specific regulation in the *SOD-1* model. Although it is known that metabolism is dysregulated in (human) ALS, the overall metabolic profile remains unknown, and we here demonstrate an essential dysregulated pathway in specific ALS (mutant) models.

Lifespan analysis of the *C9orf72* and *SOD-1* models demonstrated that mutants demonstrated a decreased lifespan compared to controls (**Figure S5C**). We supplemented PA (5 mM) at 27 °C (to introduce a mild stressor) in the *C9orf72* and *SOD-1* models; where we observed lifespan extension in the *C9orf72* and *SOD-1* G93Ac model, but not in the WTc and G85Rc model (**Figure 4H, I**). The *SOD-1* G93A strain carries GFP under the *unc-17* promotor (cholinergic neurons). We further explored neuroprotection of axons and the soma after PA supplementation, where neurons were imaged at day 8 of adulthood after 8-day PA (5 mM) exposure at 27 °C. Reconstruction of the complete neuron (including soma) demonstrated an increase in area (ratio to vehicle) in the anterior part (pharynx to vulva) (**Figure 4J**); supporting neuroprotection in this model.

To further examine functional requirement of PA and *pnk-1* we performed a paralysis assay by levamisole and RNAi of *daf-16* and *pnk-1* in the *SOD-1* model, since this model shows mutant specific dysregulated CoA metabolism. PA supplementation (5 and 10 mM) rescued paralysis in the G93Ac model. The ratio of active worms and peristaltic track length was rescued in the G93Ax strain treated with 5 mM PA, but not in the WTc and G85Rc strain (**Figure 4K**). *pnk-1* RNAi interference was more detrimental in G93Ac, and G85Rc worms compared to WTc strains by both lifespan and mobility (track length). However, *daf-16* RNAi did not have detrimental effects. These results are promising and need to be further explored, including an understanding of *daf-16* and *pnk-1* expression in other ALS models, like *TDP-43*, or other neurodegenerative disease (*e*.*g*. AD, PD) models. In our previous study, it was identified that PA levels are higher in aged/sarcopenic skeletal muscle, which could suggest a reduced requirement for CoA, increased activity of *PANK4* and ‘recycling’ of CoA, or decreased activity of *PANK1, 2* or *3*. Further exploring the PA-CoA pathway and all enzymes involved and tissue/organ specific effects could answer mechanistic questions, which can then be further translated to aging and aging related disease. To further explore PA-CoA regulation, we investigated a Chip-Seq database (**Figure S5B**) which demonstrated that *pnk-1* and *pnk-4* are targets of several genes associated to longevity. CoA is a major effector in many processes and further understanding which processes *e*.*g*. the TCA-cycle or lipid metabolism are involved in the rescue phenotype could give insight into translational directions.

## CONCLUSIONS

Our findings identify a conserved metabolic program in which the DAF-16–responsive pantothenate kinase *pnk-1* adjusts CoA biosynthesis to meet and adjust to the demands of stress and late-life physiology. By combining genetic perturbation of the PA–CoA pathway with dietary PA supplementation, our study demonstrates that boosting CoA flux is not broadly pro-longevity but instead confers selective benefits under DAF-16–associated stressors, including heat and hypoxia, and in ALS models with altered *daf-16/pnk-1* expression. The observation that PA supplementation extends lifespan and preserves cholinergic neuron integrity in these models suggests that CoA metabolism can be leveraged to modulate neurodegeneration *in vivo*. Together, these data position the pantothenate–CoA axis as a tractable metabolic node linking nutrient availability, stress-responsive transcription, and neuronal health, and provide a framework for exploring related interventions in other organisms and disease contexts where PA-CoA pathway deficits exist. These findings are nuanced and highlight the importance of understanding functional versus dysfunctional PA-CoA metabolism. We need to further understand when PA supplementation is beneficial versus detrimental and confirms a need for a better understanding and implementation of personalized medicine, where for one patient a certain medication (*e.g*. interference with the PA-CoA pathway) may be beneficial, while for another such interference by medication might be detrimental.

## Author contributions

SMJ – performed experiments and analysis, conceived and designed experiments, wrote the manuscript. AN, AB and WW performed experiments and analysis. JRT – provided supervision and expertise. PS – provided supervision and expertise, conceived and designed experiments. PAW – provided supervision, conceived, and designed experiments, wrote the manuscript. All authors read and approved of the final manuscript.

## Acknowledgments and funding

The authors would like to thank the CGC, which provided *C. elegans* strains and is funded by the NIH Office of Research Infrastructure Programs (P40 OD010440), and the *C. elegans* Reverse Genetics Core Facility, which is part of the international *C. elegans* Gene Knockout Consortium. The authors would like to thank Suny Biosciences for generating plasmids and transgenic *C. elegans* strains PWA06, PWA07 and PWA08 (https://www.sunybiotech.com/). PAW is supported by grants from Vetenskapsrådet (2022–00799), the Karolinska Institutet doctoral education program (KID), The Ulla and Ingemar Dahlberg Foundation, and Centre for Eye Research Australia philanthropic funds. PS is supported by grants from Vetenskapsrådet (2022-04418), the Swedish Brain Foundation (FO2023-0448), and the Maja & JP Åhlén Foundation.

## MATERIALS AND METHODS

### Resource and databank collection: PANK phenotypes, PANK alignment, Chip-Seq database

Phenotypes for *PANK1, -2, -3* and *-4* mutations in mice were collected from Mouse Genome Informatics (MGI GxD, https://www.informatics.jax.org/). The figures from the MGI GxD were adapted for our Supplementary Table 2. Human phenotypes of *PANK1, -2, -3* and *-4* mutations were collected on The Genome Aggregation Database (gnomAD, https://gnomad.broadinstitute.org/). Genomic sequences for the *PANK* genomes were collected on Ensembl (https://www.ensembl.org/) and protein sequences on NCBI (https://www.ncbi.nlm.nih.gov/). Functional domains in protein sequences were identified by SMART (https://smart.embl.de/). Nucleotide and amino acid sequence alignments were performed with MEGA12 (https://www.megasoftware.net/) and ClustalW alignments. The Chip-Seq database Chip-Atlas (https://chip-atlas.org/peak_browser) was used to identify genes that have *pnk-1* or *pnk-4* as a target gene via the ‘Target gene’ function searching for +/-5 KB from the transcription starting site.

### C. elegans strains and maintenance

Worm strains and *E. coli* OP50 were obtained from the Caenorhabditis Genetics Centre (CGC, https://cgc.umn.edu/). Worms were maintained on 6 cm dishes at 15°C or 20°C, depending on the strain, on standard nematode growth medium (NGM) with *E. coli* OP50 10x concentrated. The strains used in this study are described in **Supp. Table 7**. Chemicals used in the assays are listed in **Supp. Table 8**.

### Overexpression of pnk-1 and pnk-4

Overexpression lines of *pnk-1, pnk-4, Y65B4A*.*8* and *pnk-1::-4* were produced by Suny Biosciences. cDNA sequences were collected on Ensembl. Plasmids in the pPD95.77 vector backbone (including a GFP tag) were constructed with a *rpl-28* promoter, followed by the cDNA sequence of the gene of interest and followed by an *unc-54* 3’UTR. This resulted in the four following strains; PWA06 (Ex[Prpl-28-pnk-4 cDNA-GFP-unc-54 3’UTR]), PWA07 (Ex [Prpl-28-pnk-1 cDNA-SL2-pnk-4 cDNA-GFP-unc-54 3’UTR]), PWA08 (Ex [Prpl-28-Y65B4A.8 cDNA-GFP-unc-54 3’UTR]). Injection concentrations were 10 ng/µl for PWA06 and PWA08, and 2 ng/µl for PWA07 (due to failure of generating lines with 10 ng/µl). Generation of PWA05 (overexpression of *pnk-1*) failed with injecting 10 and 2 ng/µl of plasmid.

### Longevity and healthspan

Longevity and healthspan assays were performed on 6 cm dishes with NGM supplemented with 50 µM FuDR to prevent the generation of offspring. Supplementation of PA or CoA was performed by seeding of diluted PA or CoA (from 25 mM stocks stored at -20°C) in 1% PFA treated *E. coli* OP50 (1 hour treatment followed by 4x washes and 10x concentration). We used 5 mM PA and 0.5 mM CoA, a dose within the range commonly used for dietary metabolite supplementation in *C. elegans* and previously shown to modulate lifespan and stress responses without overt toxicity. This concentration was selected to reliably engage the PA–CoA pathway in vivo while avoiding developmental delays or gross behavioral defects. L4 to young adults (recognized by a white vulval spot) were moved to supplemented or control plates which represents day 1 of adulthood in all experiments. A minimum of 10 and maximum of 30 worms were kept per plate and multiple plates were used in 2 to 3 individual experiments. Re-supplementation was performed by adding 20 µL of fresh diluted supplement onto plates every three days until day 16 of adulthood or moving worms to freshly seeded plates. Videos of worms were recorded (Wormlab, MBF Bioscience) on day 3, 5, 8 and 11 of adulthood for supplementation studies. Worms were recorded every 2-3 days for life, dead, censored, or lost status; censored worms showed signs of internal hatching or intestine leakage. Depending on the strain the experiments were performed at 15°C or 20°C.

### RNA interference

*E. coli* HT115(DE3) clones containing dsDNA were obtained from the Julie Ahringer library. Clones containing dsDNA for *pnk-1* (C10G11.5) and *pnk-4* (C42D8.3), were grown in liquid broth + 25 µg/ml ampicillin and colonies were 4x concentrated before seeding 100 µL per 1 mM IPTG (+ 25 µg/ml carbenicillin) plates. L3 wild-type (N2), daf-2 (*e1370*), daf-16 (*mgdf50*) or daf-2::daf-16 (*mgdf50::m65*) worms were placed on plates with HT115(DE3) clones of interest (with *L4440* as a (empty) vehicle and *pos-1* as positive control). The L4 stage F2 generation was used for experimental plates where worms were moved to fresh plates every 4 days until there was no more reproduction. On day 4 and 6 of adulthood, videos of worms were recorded (Wormlab) and lifespan was recorded every 2-3 days. Depending on the strain the experiments were performed at 15° C or 20° C.

### Stress assays

Standard NGM + FuDR plates with 5 mM PA or 0.5 mM CoA in 1% PFA *E. coli* OP50 were prepared as described above under ‘*Longevity and healthspan’*. For oxidative stress assays, worms were moved on day 1 of adulthood to plates with an additional supplement of 1.5 µM rotenone dissolved in DMSO in the OP50 and moved to fresh plates on day 8 of adulthood, at 20 °C. For hypoxic stress assays, worms were moved on day 1 of adulthood to plates with an additional supplement of 2 mM sodium-azide in OP50 and moved to fresh plates with 4 mM sodium-azide on day 4 of adulthood, at 20 °C. For heat stress assays, worms were moved to supplemented plates on day 1 of adulthood and moved to 30°C until death (no re-supplementation performed, alive, dead, censored status was scored every day). Generally, during ‘stress’ experiments, the worms were scored for alive, dead, lost or censored status every 1-3 days.

### Paralysis assays

Standard NGM + FuDR plates with 5 mM or 10 mM PA in 1% PFA *E. coli* OP50 were prepared as described above under ‘Longevity and healthspan’. ALS SOD-1 worms were moved to PA plates on day 1 of adulthood (L4 stage) and exposed to 27° C until day 3 of adulthood. On day 3 of adulthood, standard NGM plates -FuDR and 100 µM levamisole diluted in OP50 were prepared and 10 worms were moved to levamisole+ plates. Spontaneous movement by crawling (no tapping of plates or tapping of worms with a worm pick) was counted every 15 minutes for the first hour followed by every 30 or 60 minutes afterwards for the next 3 hours. At 60 minutes exposure, mobility was measured with the Wormlab system.

### CoA quantification (luminescence)

Whole body *C. elegans* coenzyme A levels were measured using the Coenzyme A Assay Kit Fluorometric Green kit (Abcam, ab138889). Briefly, L4 worms were moved to standard NGM plates seeded with OP50. For temporal measurements, worms were aged according to the day (1, 3, 5, 8, 11 or 15 of adulthood) while being moved every ∼3 days to get rid of offspring. For OE, ALS and KO measurements, worms were harvested on day 3 of adulthood. On the day of harvest, 5 to 10 worms were lysed via sonication in 225 µL of distilled H_2_O. 100 µL of medium was used for the CoA kit (50 µL in duplicate) and 100 µL (50 µL in duplicate) for a protein determination assay (Bradford, Thermo Fisher); CoA levels were presented in µg CoA per µg of protein.

### RT-qPCR

For mRNA analysis, either ∼10 worms (adults) were lysed in 3 µL lysis buffer (5 mM Tris pH 8.0, 0.5% Triton-X, 0.5% Tween20, 0.25 mM EDTA, 1 mg/ml Proteinase K) in the thermocycler (65° C for 10 min followed by inactivation of Proteinase K for 1 min at 85° C) or lysed samples used for CoA quantification (see above) were directly used (3 µL). RT was performed by using iScript (*BioRad*) in a total volume of 10 or 20 µL followed by a qPCR in a total volume of 10 µL for the following genes; *pnk-1* (*fwd: 5’-cgtgattctaccagcatcgg-3’, rev: 5’-acgatcagtccacacctctt-3’*), *pnk-4* (*fwd: 5’-ccgtgggttactagctggaa-3’, rev: 5’-gccaactccaccatttctcg-3’*), *daf-16* (*fwd: 5’-gctcccttcactcgacactt-3’, rev: 5’-ttgagttcgggaacggaaag-3’*), *mtl-1* (*fwd: 5’-ggcttgcaagtgtgactgc-3’, rev: 5’-tttccgcacttgcattgctt-3’*), *hsp-16*.*2 (fwd: 5’-tccatctgagtcttctgagattgtt-3’, rev: 5’-tgagacgttgagattgatggca-3’)* (*mtl-1* and *hsp-16*.*2* by Zhang et al. (Zhang et al., 2024) and *pmp-3, cdc-42, tba-1* as published by Hoogewijs et al. (Hoogewijs et al., 2008).

### Imaging and analysis

*C. elegans* reporter strains were moved to CoA and PA supplemented plates as described under ‘Longevity and healthspan’ on day 1 of adulthood and re-supplemented on day 4 and 7 (by adding 20 µL of fresh supplement or by moving worms to fresh plates). On day 5 and 10 of adulthood, worms were moved to a 2% agar pad on a glass slide and anesthetized in levamisole hydrochloride and worms were imaged with a Leica Stellaris 5X. SJ4103 and SJZ42 were imaged with a 63x oil objective; RW1596 was imaged with a 40x glycerol objective; and VS20 was imaged with a 20x glycerol objective. SJ4103 was analyzed with the MitoAnalyzer plugin in FIJI; RW1596 was analyzed with the MuscleMetrics plugin in FIJI; SJZ42 was analyzed in Imaris, where objects were created of particles and fluorescence intensity was measured inside created objects. VS20 was analyzed in FIJI via thresholding, after which a selection of the threshold was created, and fluorescence intensity was measured inside the selection.

### Data analysis, statistics, and availability

A Wilcoxon rank test was used to compare groups. *C. elegans* lifespan data was analyzed by Kaplan-Meier with a *p-value* calculation by Log-rank (see lifespan data in **Supp. Dataset 2**). Lifespan data was right censored with a death event scored as 1 and lost or censored worms scored as 0. All statistical analysis and graphics were performed in R. All generated data is available within this manuscript and the associated supplementary files; any additional protocols or code are available on request.

