## Supplementary Figures and Tables for "Supplementation via DAF-16 and *pnk-1* driven pantothenate–coenzyme A flux improves disease related stress resistance in *C. elegans*"

**Supplementary Materials**

The following supplementary materials are attached:

Supplementary Figures 1-5

Supplementary Tables 1-8.

Supplementary Datasets 1 and 2 are available as .csv

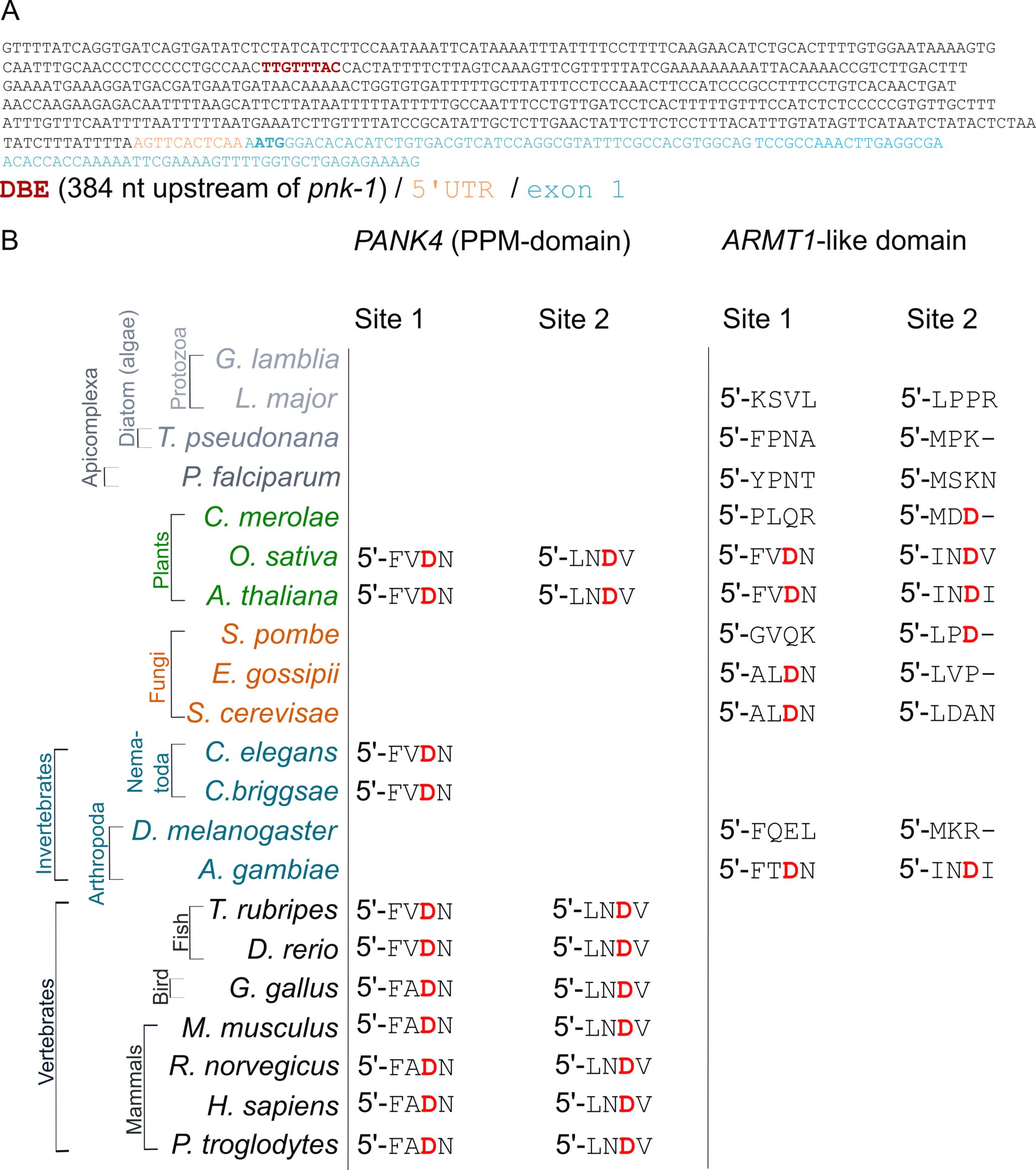
**Supplementary Figure 1. DAF-16 binding element in *pnk-1* and functional alignment of PPM in PANK4.** (**A**) Representation of the DAF-16 Binding Element (**DBE**; 5’-TTGTTTAC-3’) in the genomic sequence of *C. elegans pnk-1.* (**B**) Functional alignment of metal dependent protein phosphatase (PPM) in *PANK4* and the separate *ARMT-1* (Acidic Residue Methyltransferase 1, metal-dependent phosphatase) domain in different organisms.

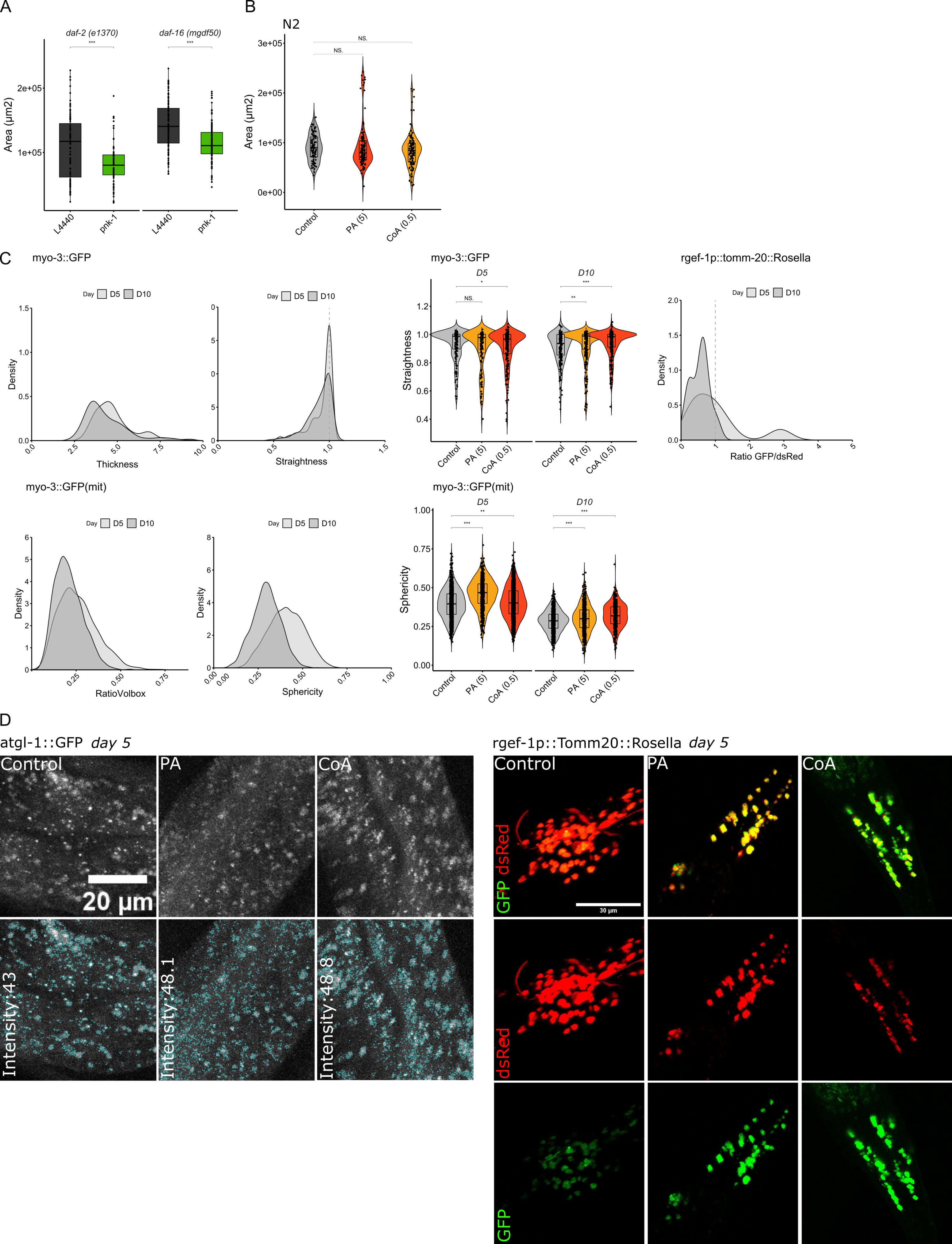
**Supplementary Figure 2. Additional: systemic effects of aging and PA supplementation by reporter strains.** (**A**) Growth phenotype of *daf-2 (e1370)* and *daf-16 (mgdf50)* mutants exposed to *pnk-1* RNAi measured at day 4-6 of adulthood. (**B**) Growth phenotype of wild type N2 worms (at day 5 of adulthood) supplemented with 5 mM PA or 0.5 mM CoA. (**C**) Density plots of quantified numbers of *C. elegans* reporter strains including myo-3::GFP, myo-3::GFP(mit) and rgef-1p::tomm29::Rosella, demonstrating aging trends at day 5 to day 10 of adulthood. These trends represent aging phenotypes of muscle, mitochondria and mitophagy; where at day 10 density for all parameters is shifted to the left, representing *e.g.*  decreased straightness of myosin and decreased sphericity of mitochondria. (**D**) Images representing atgl-1::GFP worms supplemented with 5 mM PA or 0.5 mM CoA (top row) and in blue (bottom row) quantification of thresholded GFP signal. Images representing rgef-1p::Tomm20::Rosella worms supplemented with 5 mM PA or 0.5 mM CoA where a GFP/dsRED merge is visualized in the top row, and separate channels for dsRED and GFP are shown below. *p = * < 0.05, ** < 0.01, *** < 0.001, **** < 0.0001.

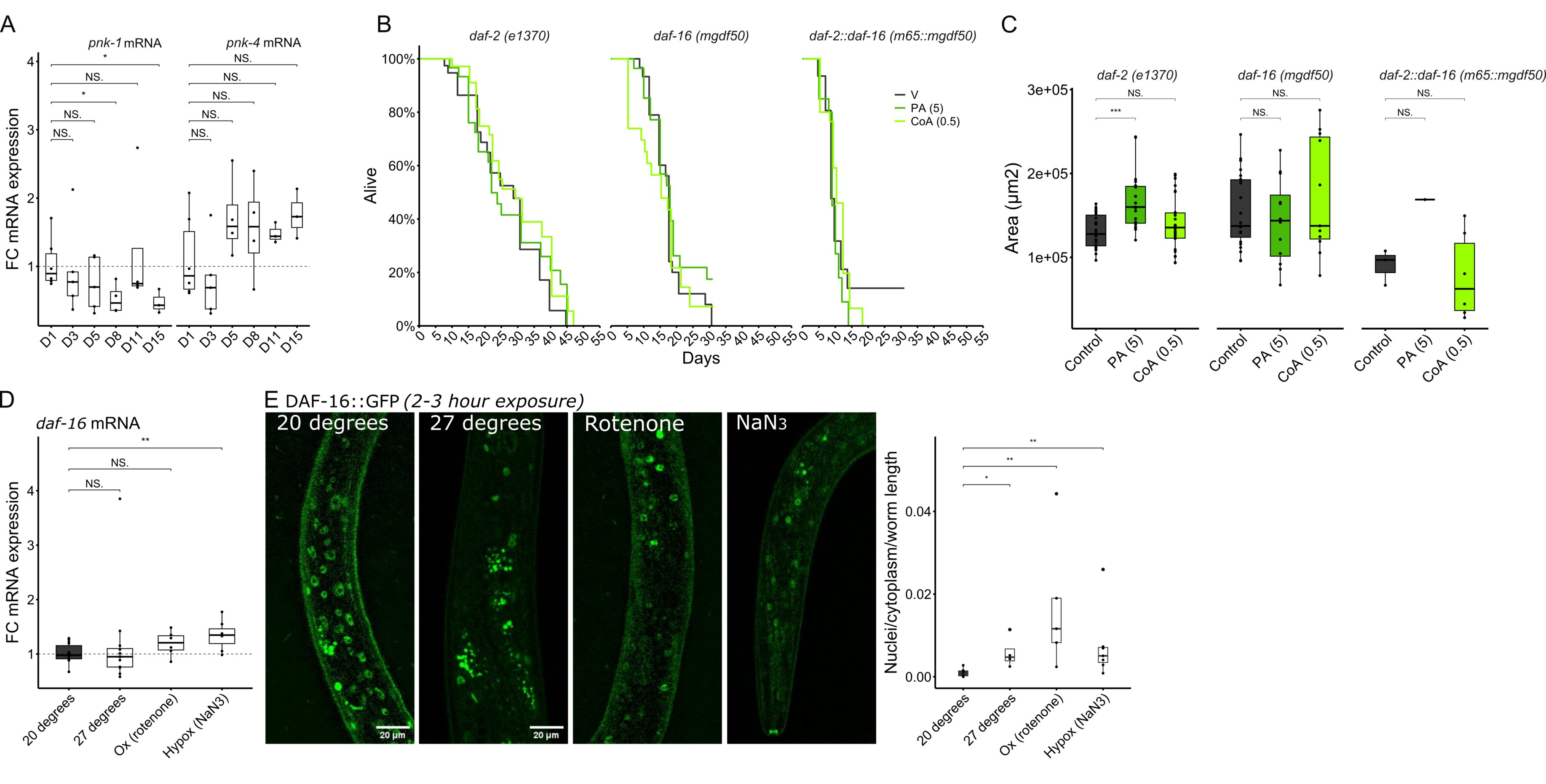

**Supplementary Figure 3. Additional: *pnk-1* is essential in wild-type worms, regulated via *daf-16*, and PA supplementation improves DAF-16 associated stress resistance.** (**A**) Temporal *pnk-1* and *pnk-4* mRNA expression by RT-qPCR, which suggests increased expression of *pnk-4* later in life, but results are variable. (**B**, **C**) Lifespan by Kaplan-Meier and growth phenotype after PA (5 mM) and CoA (0.5 mM) supplementation in *daf-2 (e1370)* and *-16 (mgfdf50)* mutant worms. (**D**) *daf-16* mRNA expression by RT-qPCR upon stress exposure. (**E**) DAF-16 translocation upon stress exposure, where translocation in nuclei per DAF-16 in cytoplasm per worm length was plotted. *p = * < 0.05, ** < 0.01, *** < 0.001, **** < 0.0001.

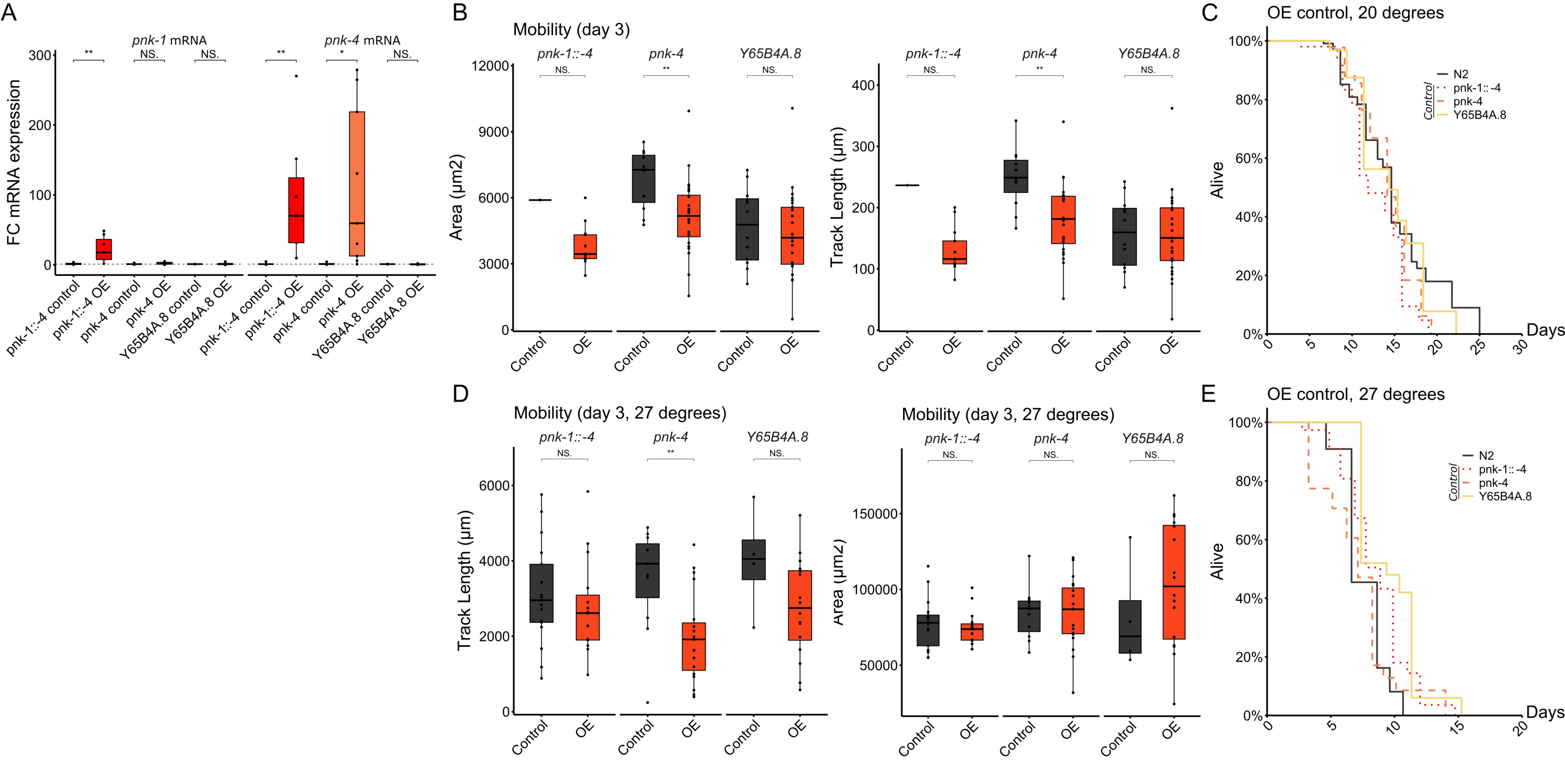
**Supplementary Figure 4. Additional: OE strains and longevity phenotypes**. (**A**) mRNA expression by RT-qPCR of *pnk-1* and *-4* in overexpression strains of *pnk-1::-4, pnk-4* and *Y65B4A.8* compared to their respective control strain (the respective control strain has a N2 background). (**B, D**) Worm growth and mobility of OE lines at day 3 of adulthood at 20° C and 27° C. (**C, E**) Lifespan by Kaplan-Meier of the respective control strains of the generated OE strains (the respective control strain has a N2 background) at 20° C and 27° C. *p = * < 0.05, ** < 0.01, *** < 0.001, **** < 0.0001.

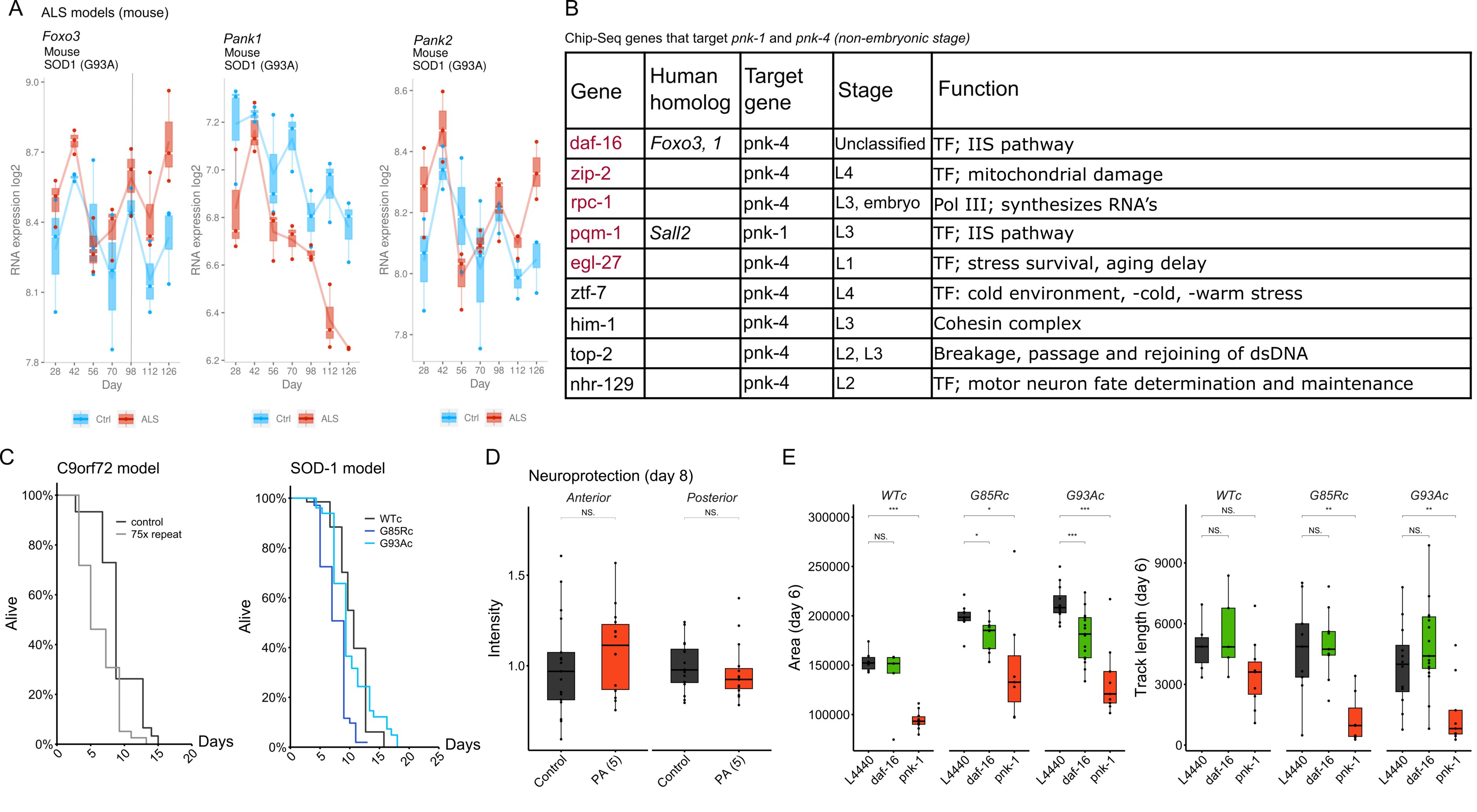
**Supplementary Figure 5. Additional: ALS cell atlas, Chip-Seq targets and ALS phenotypes.** (**A**) RNA expression levels of *Foxo3,* *Pank1* and *Pank2*, identified on ALS cell atlas (https://alscellatlas.org/ (Månberg et al., 2021)) over time (126 days); in blue: RNA expression levels of control mice, in red: RNA expression levels of *SOD-1* G93A mice. (**B**) Chip-Seq identified transcription factors that target *pnk-1* or *-4* in stages older than L3 (identified at https://chip-atlas.org/peak_browser), in red text: genes that are associated to longevity in *C. elegans*. (**C**) Lifespan of two ALS *C. elegans* models; a *C9orf72* model and *SOD-1* model, where mutants demonstrate a decreased lifespan as published in original papers. (**D**) Intensity of GFP in reconstructed cholinergic neurons in the *SOD-1* G93Ac strain. (**E**) Growth phenotype and mobility at day 6 of adulthood of ALS *SOD-1* worms exposed to *daf-16* and *pnk-1* RNAi. *p = * < 0.05, ** < 0.01, *** < 0.001, **** < 0.0001.

**Supplementary Table 1.** Associated phenotypes and SNPs in pantothenate kinases (human and mouse).

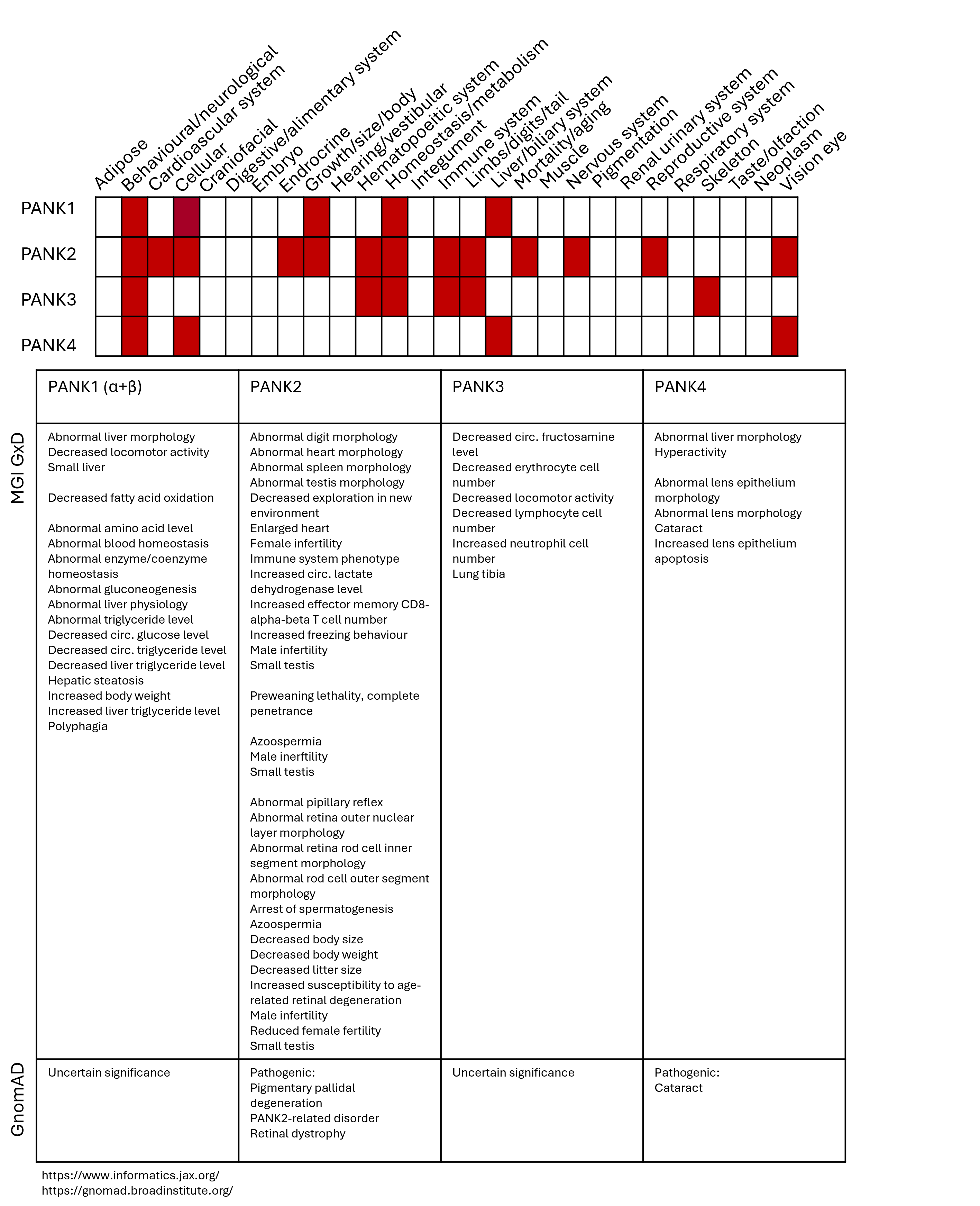

**Supplementary Table 2.** Observations on PANK

| Gene | Finding | Year | Reference |
| --- | --- | --- | --- |
| PANK1, mouse  PANK1β, mouse | Increased PANK1 expression increased CoA and depleted the PA pool  Stimulated by CoA and inhibited by acetyl-CoA | 2000 | Rock et al. |
| PANK3, | Vit. B9 activated, and c-Myc repressed PANK3 | 2025 | Zhang et al. |
| PANK4, human  PANK4, human  PANK4, mouse | A pseudo-pantothenate kinase  Has a functional phosphatase domain  Whole body knock-out decreased body weight, tissue mass and length in females, but not males | 2019  2022  2025 | Yao et al.  Dibble et al.  Miranda-Cervantes et al. |

**Supplementary Table 3.** Protein homology of *C. elegans* proteins to *human* proteins.

| Gene | Uniprot ID | Alphafold hits  (*Homo sapiens*) | Length (AA) | % protein homology  (to human) | Conserved Domains  (BLAST, CD) |
| --- | --- | --- | --- | --- | --- |
| *PANK4* | Q9NVE7 |  | 773 |  | PanK-II_Pank4 (36-366)  PLN02902 (17-771) |
| *pnk-1* | H2KYP7_CAEEL | *PANK4* | 449 | 67% (Alphafold) | PanK-II_Pank4 (115-441) |
| *pnk-4* | Q18580_CAEEL | *PANK4* | 755 | 59% (Alphafold) | PLN02902 (104-748) |
| *PPCS* | Q9HAB8 |  | 311 |  | PRK05579 (15-302) |
| *Y71H2AM.6* | Q9BL36 | *PPCS* | 277 | 44% (Alphafold) | PRK05579 (20-268) |
| *PPCDC* | Q96CD2 |  | 204 |  | PLN02496 (16-204) |
| *F25H9.6* | P91988_CAEEL | *PPCDC* | 237 | 68% (Alphafold) | PLN02496 (42-222) |
| *COASY* | Q13057 |  | 564 |  | cd02164 (193-339)  cd02022 (360-536) |
| *Y65B4A.8* | Q9BL56_CAEEL | *COASY* | 461 | 60% (Alphafold) | cd02164 (102-245)  cd02022 (267-444) |
| *Foxo3* | O43524 |  | 673 |  | cd20061 (157-239)  pfam16675 (432-511)  pfam16676 (606-644) |
| *daf-16* | O16850 | *FoxO3, FoxO1* | 541 | 66%, 57% (Alphafold) | cd20032 (176-255) |

**Supplementary Table 4.** DBE conservation *Caenorhabditis* and other species in *pnk-1* and pantothenate kinases *(max 5000 nt upstream of coding region start codon / ATG)*

| Species | Gene | DBE | Location |
| --- | --- | --- | --- |
| *Caenorhabditis briggsae* | *pnk-1* | TTGTTTAC | 384 upstream ATG |
| *C. remanei* | *pnk-1* | TTGTTTAC | 390 upstream ATG |
| *C. brenneri* | *pnk-1* | TTGTTTAC | 442 upstream ATG |
| *C. elegans* | *pnk-1* | TTGTTTAC | 396 upstream ATG |
| *C. japonica* | *pnk-1* | TTGTTTAC | 1787 upstream ATG |
| *Drosophila melanogaster* | *fumble* | TTGTTTAC | 629 upstream ATG |
| *Mus musculus* | *Pank1* | TGTTTAC | 592 upstream ATG |
|  | *Pank2* | TTGTTTAC | 1592 upstream CTG |
| *Homo sapiens* | *Pank1* | TGTTTAC | 2087 upstream ATG |

**Supplementary Table 5.** Mutations in *C. elegans* knockout strains (in order of the PA-to-CoA pathway).

| Gene | Strain (CGC) | Location (chromosome) | Genotype | Deletion |
| --- | --- | --- | --- | --- |
| *pnk-1* (C10G11.5) | VC927 | I | pnk-1(ok1435) I/hT2 [bli-4(e937) let-?(q782) qIs48] (I;III). | ok1435: 773 bp  *Intron 2 to exon 5* |
| *pnk-4* (C42D8.3) | RB1527 | X | pnk-4(ok1832) X. | ok1832: 1034 bp  *Intron 3 to exon 5* |
| *Y71H2AM.6* | VH7172 | III | Y71H2AM.6(hd7172[LoxP + myo-2p::GFP::unc-54 3' UTR + rps-27p::neoR::unc-54 3' UTR + LoxP]) III. | 911 bp  *-90 bp (from 5’UTR) to intron 3* |
| *F25H9.6* | RG3297 | V | +/nT1[umnIs49] IV; F25H9.6(ve797[LoxP + myo-2p::GFP::unc-54 3' UTR + rps-27p::neoR::unc-54 3' UTR + LoxP])/nT1 V. | 1068 bp  *Exon 1 to exon 4* |
| *Y65B4A.8* | VC4781 | I | Y65B4A.8(gk5849[loxP + myo-2p::GFP::unc-54 3' UTR + rps-27p::neoR::unc-54 3' UTR + loxP])/+ I. | 3386 bp  *-17 bp (from 5’UTR) to exon 4* |

**Supplementary Table 6.** *C. elegans* transgenic ALS models.

| Model | Mutation | Strain (CGC) | Allele | Method |
| --- | --- | --- | --- | --- |
| SOD-1 | WTc  G85Rc  G93Ac | HA2986  HA3299  HA2987 | sod-1(rt448[sod-1WTC]) II.  sod-1(rt451[sod-1(G85RC)]) II.  sod-1(rt449[G93AC]) II; vsIs48. | CRISPR/Cas9 |
| C9orf72 | Control  75 GGGGCC repeats | KRA317  KRA315 | kasIs9 [snb-1p::(delta)C9 ubi + myo-2::GFP]  kasIs7 [snb-1p::C9 ubi + myo-2::GFP]. | Microinjection |

**Supplementary Table 7.** List of strains used.

| Strain | Genotype |
| --- | --- |
| Wild-type N2 | wild type isolate |
| CB1370 | daf-2(e1370) III |
| CF1139 | daf-16(mu86) I; muIs61(pKL78) daf16::GFP + rol-6(su1006) |
| GA158 | daf-16(mgDf50) I; daf-2(m65) III |
| GR1307 | daf-16(mgDf50) I |
| HA2986 | sod-1(rt448[sod-1WTC]) II. |
| HA2987 | sod-1(rt449[G93AC]) II; vsIs48. |
| HA3299 | sod-1(rt451[sod-1(G85RC)]) II. |
| KRA315 | kasIs7 [snb-1p::C9 ubi + myo-2::GFP]. |
| KRA317 | kasIs9 [snb-1p::(delta)C9 ubi + myo-2::GFP] |
| PWA06 | Prpl-28-pnk-4 cDNA-GFP-unc-54 3’UTR |
| PWA07 | Prpl-28-pnk-1 cDNA-SL2-pnk-4 cDNA-GFP-unc-54 3’UTR |
| PWA08 | Prpl-28-Y65B4A.8 cDNA-GFP-unc-54 3’UTR |
| RB1527 | pnk-4(ok1832) X), VH7172 (Y71H2AM.6(hd7172[LoxP  + myo-2p::GFP::unc-54 3' UTR  + rps-27p::neoR::unc-54 3' UTR + LoxP]) III |
| RG3297 | ((+/nT1[umnIs49] IV; F25H9.6(ve797[LoxP  + myo-2p::GFP::unc-54 3' UTR  + rps-27p::neoR::unc-54 3' UTR  + LoxP])/nT1 V |
| RW1596 | stEx30 [myo-3p::GFP::myo-3  + rol-6(su1006) |
| SJ4103 | (zcIs14 [myo-3::GFP(mit)) |
| SJZ42 | foxEx3 [rgef-1p::tomm-20::Rosella |
| VC4781U | Y65B4A.8(gk5849[loxP  + myo-2p::GFP::unc-54 3' UTR  + rps-27p::neoR::unc-54 3' UTR + loxP])/+ I) |
| VC927 | pnk-1(ok1435) I/hT2 [bli-4(e937) let-?(q782) qIs48] (I;III) |
| VS20 | hjIs67 [atgl-1p::atgl-1::GFP + mec-7::RFP |

**Supplementary Table 8.** List of chemicals used in experiments.

| Chemical | Supplier | Catalog # |
| --- | --- | --- |
| Agar | Sigma Aldrich | 05039-500G |
| Bacto-peptone | Sigma Aldrich | P6838-500G |
| Calcium Chloride | Sigma Aldrich | C1016-100G |
| Cholesterol  Coenzyme A | Sigma Aldrich  Sigma Aldrich | C8667-5G  ATE514323213 |
| Dimethyl sulfoxide | Sigma Aldrich | D2438-5X10ML |
| 5-Fluorodeoxyuridine | Sigma Aldrich | F0503-100MG |
| KH2PO4 | Sigma Aldrich | P5655-1KG |
| K2HPO4 | Sigma Aldrich | P3786-1KG |
| Levamisole hydrochloride | Sigma Aldrich | 31742-250MG |
| Liquid broth | Gibco | 10855-001 |
| M9 | Sigma Aldrich | M6030-1KG |
| Pantothenic acid | Sigma Aldrich | P5155-100G |
| Phosphate buffered saline  Rotenone | Gibco  Sigma Aldrich | 70011-044  R8875 |
| Sarcosine | Sigma Aldrich | 131776-100G |
| Sodium chloride  Sodium-azide | Sigma Aldrich  Sigma Aldrich | S5886-500G  S2002 |
